# Higher-order epistatic networks underlie the evolutionary fitness landscape of a xenobiotic-degrading enzyme

**DOI:** 10.1101/504811

**Authors:** Gloria Yang, Dave W Anderson, Florian Baier, Elias Dohmen, Nansook Hong, Paul D Carr, Shina Caroline Lynn Kamerlin, Colin J Jackson, Erich Bornberg-Bauer, Nobuhiko Tokuriki

**Author notes:** To whom correspondence should be addressed: Nobuhiko Tokuriki, Ph. D., Michael Smith Laboratories, University of British Columbia, Vancouver, V6T 1Z4, BC, Canada.

## Abstract

Characterizing the adaptive landscapes that encompass the emergence of novel enzyme functions can provide molecular insights into both enzymatic and evolutionary mechanisms. Here, we combine ancestral protein reconstruction with biochemical, structural, and mutational analyses to characterize the functional evolution of methyl-parathion hydrolase (MPH), a xenobiotic organophosphate-degrading enzyme. We identify five mutations that are necessary and sufficient for the evolution of MPH from an ancestral dihydrocoumarin hydrolase. In-depth analyses of the adaptive landscapes encompassing this evolutionary transition revealed that a complex interaction network, defined in part by higher-order epistasis, determined the adaptive pathways that were available. By also characterizing the adaptive landscapes in terms of their functional activity towards three other OP substrates, we reveal that subtle differences in substrate substituents drastically alter the enzyme’s epistatic network by changing its intramolecular interactions. Our work suggests that the mutations function collectively to enable substrate recognition via subtle structural repositioning.

## Introduction

How evolution generated a tremendous repertoire of diverse enzymatic functions is one of the central questions in evolutionary biochemistry. Understanding the molecular mechanisms that underlie the evolution of new enzyme functions requires 1) the identification of a minimal set of genotypic changes that are necessary and sufficient to cause the emergence and optimization of a new function, and 2) an understanding of the biophysical molecular changes caused by these mutations that resulted in a novel function. Recent advances in experimental evolution and phylogenetically informed ancestral sequence reconstruction (ASR) aid in such endeavours by unveiling a set of adaptive mutations that allow substantial functional transitions^1–9^. Further biochemical, biophysical and structural characterization of enzyme variants enables us to uncover the molecular basis of the functional transition. Elucidating the molecular role of each genotypic change, however, is often difficult due to the prevalence and complexity of epistasis, *i.e*., the phenomenon in which the effect of a mutation varies significantly depending on the presence or absence of other mutation(s)^10–15^. Consequently, the mutational effect observed in a particular genetic background, *i.e*., the wild-type genotype, may not always accurately reflect the impact that the particular mutation had during the actual historical evolution if the mutation was acquired after other earlier mutations had already fixed. Understanding the molecular basis of epistasis is thus essential if we are to effectively investigate the sequence-structure-function relationships that govern how proteins evolve novel functions.

Adaptive fitness landscapes, which can be determined by generating and functionally assaying all possible combinations of the mutations responsible for a new function, have become a powerful tool for studying the evolutionary and biophysical origins of novel functions by unveiling the potential adaptive pathways that connect the ancestral and derived genotypes^10–12,14,16–18^. Recently developed statistical methods enable us to assess and quantify the degree of epistasis, including high-order interactions (those that involve interactions between more than two mutations), which provide a comprehensive view of epistasis and the dominant interactions that drive it^19–22^. Moreover, because epistasis reflects interactions between amino acid changes, adaptive landscapes also provide critical insight into the underlying molecular interactions both within an enzyme and between enzyme and substrate. With this study, we integrate an analysis of adaptive landscapes with a comprehensive assessment of the intra-and intermolecular epistatic interactions underlying the evolution of a novel organophosphate hydrolase (OPH) enzyme in order to obtain a detailed view of how enzymes can evolve novel catalytic functions.

In response to the industrial synthesis and application of organophosphate (OPs) pesticides in the early 20^th^ century, multiple classes of OPH enzymes have been isolated from various strains of soil bacteria, where they confer a selective advantage to the bacteria by allowing OPs to be used as a source of phosphate and carbon^23^. One example is methyl-parathion hydrolase (MPH), which was isolated from *Pseudomonas* sp. WBC-3 in the polluted soil near a methyl-parathion production factory^24^. Because the selection pressure is relatively simple and well understood, this enzyme provides an excellent opportunity to investigate how evolution produces novel catalytic functions. Presently, however, the evolutionary and molecular mechanisms underlying the evolution of MPH are largely unknown.

Here, we combine ASR with a robust functional, structural, and mutational characterization of the ancestral and derived enzymes to identify a minimal historical set of five genotypic changes that were responsible for optimizing the novel enzymatic function. We subsequently characterized the adaptive landscape that encompasses this evolutionary transition as defined by these five key historical substitutions for activity against four different OP substrates. We then develop and apply extensive statistical analyses to understand how epistatic interactions defined this landscape for each substrate. In particular, we unveil the complex interaction network that underlies the fine-tuning of enzyme specificity for methyl-parathion activity over other similar OP compounds.

## Results

### MPH evolved from a dihydrocoumarin hydrolase enzyme

Previous studies have found that MPH is closely related to a group of enzymes that exhibit high hydrolytic activity towards the lactone dihydrocoumarin (DHC), which is an intermediate compound in the degradation pathways of a number of aromatic hydrocarbons in bacteria^25–27^(**Fig. 1** and **Supplementary Fig. 1**). In order to rigorously establish the evolutionary relationship between DHC hydrolases (DHCH) and MPH enzymes, we performed multiple sequence alignment and phylogenetic analyses on 153 publicly available sequences that are most closely homologous to *Pseudomonas* sp WBC-3 (MPH-wt) (**Fig. 1** and **Supplementary Fig. 2)**. The vast majority of these sequences (all but six) exhibit a maximum of ∼60% amino acid sequence identity with MPH-wt; the remaining six sequences form a small clade in which they share >90% amino acid sequence identity. We then inferred the maximum likelihood amino acid sequence for the ancestral phylogenetic nodes that separate the MPH-like clade from the paraphyletic DHCH groups (**Supplementary Fig. 2**). AncDHCH1 refers to the phylogenetic node that is the common ancestor for the MPH clade and a few DHCHs, including (*Js*DHCH) from *Janthinobacterium* sp. HH01. AncDHCH2 and AncDHCH3 represent deeper phylogenetic nodes that are ancestral to AncDHCH1 and to other DHCHs (**Fig. 1**). The prediction of each ancestral sequence reconstruction receives a high level of statistical support, with average posterior probabilities >0.89 (**Supplementary Fig. 3** and **Supplementary Table 1**). Overall, AncDHCH1 has 89% amino acid sequence identity to MPH-wt (differing at 31 amino acid substitutions and a single amino-acid insertion), and 73% amino acid sequence identity to *Js*DHCH (72 substitutions).

**Figure 1.**
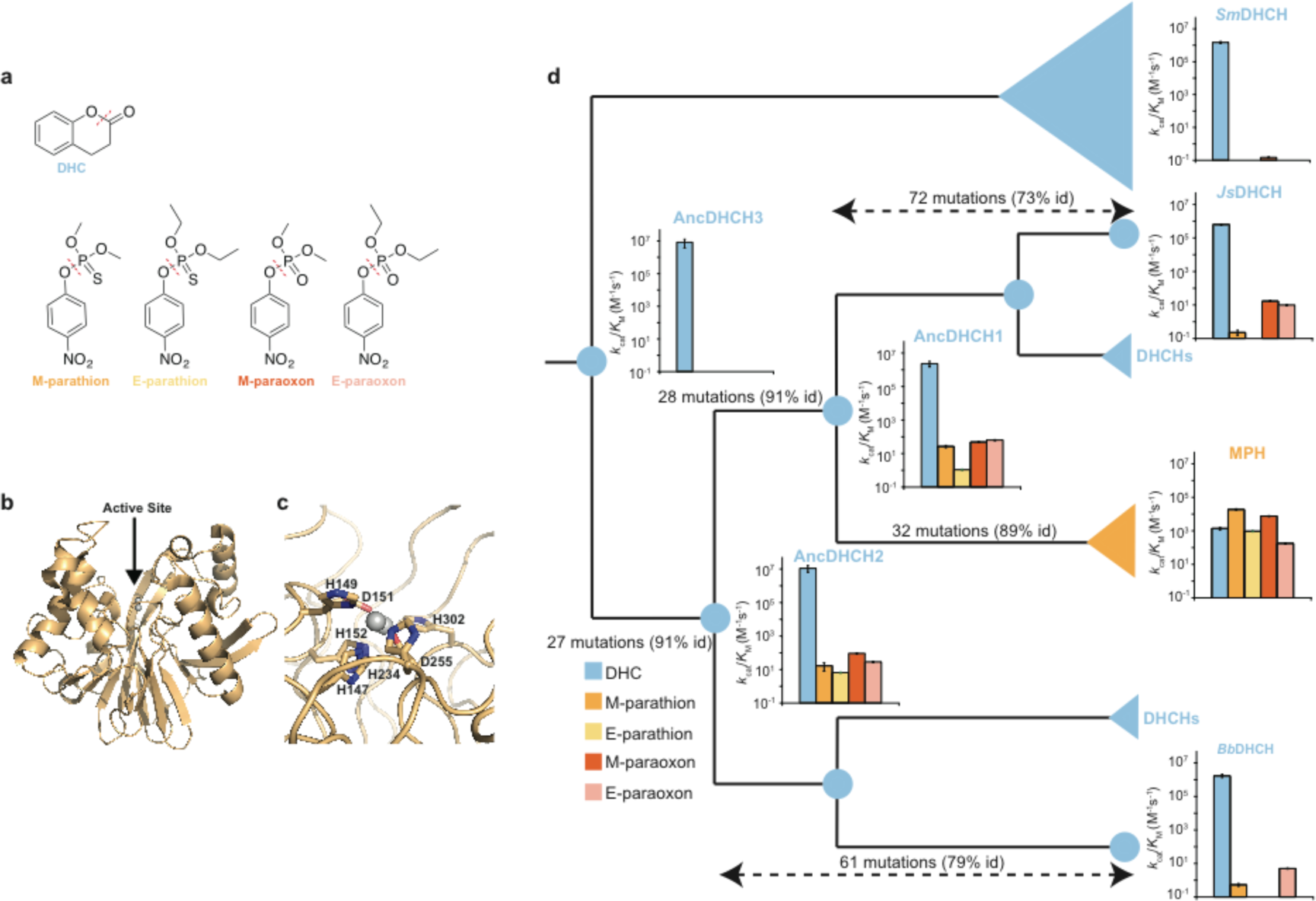
Phylogeny and phenotype of methyl-parathion hydrolase (MPH) **a**, Chemical structures of the four organophosphate (OP) and dihydrocoumarin (DHC) substrates utilized in this study. Dashed red lines indicate the bond that is cleaved during the reaction. Proposed mechanisms of MPH for OP and DHC hydrolase activities are described in **Supplementary Fig. 5**. **b**, Cartoon representation of the crystal structure MPH from *Pseudomonas* sp. WBC-3 (PDB entry 1P9E). The two active site metal ions are shown as grey spheres. **c**, Cartoon representation of the active site of MPH. The active site metal ions are shown as grey spheres. Residues coordinating the metals are highlighted as sticks. **d**, Schematic presentation of the phylogeny of MPHs, DHCHs and predicted ancestral sequences. A full phylogeny of MPH is described in **Supplementary Fig. 2**. Spheres represent individual sequences while wedges depict groups of sequences. The number of mutations and the level of identity between the sequences are indicated on the branches. The kinetic activities of the characterized sequences for DHC and four OP compounds are shown next to their corresponding nodes on the phylogeny. A full description of the kinetic parameters of the sequences can be found in **Supplementary Table 2**.

To narrow our analysis to the specific lineage on which MPH functionality evolved, we synthesized, expressed, and purified four extant enzymes (MPH-wt, *Js*DHCH, *Bb*DHCH and *Sm*DHCH) and three reconstructed ancestral enzymes (AncDHCH1, AncDHCH2, AncDHCH3). Each enzyme was characterized for their hydrolytic activity towards DHC (the hypothetical ancestral substrate), methyl-parathion (the canonical MPH OP substrate) as well as three additional OP compounds (ethyl-parathion, methyl-paraoxon and ethyl-paraoxon) (**Fig. 1** and **Supplementary Table 2**). All four OP substrates contain the same leaving group (*p*-nitrophenol) but vary in terms of their substituent groups (*i.e.*, sulphur *vs.* oxygen and ethyl *vs.* methyl). The three extant DHCH homologs and all ancestral enzymes exhibit high DHCH activity (*k*_cat_/*K*_M_ ≥ 10^6^ M^-1^s^-1^) and relatively low OP hydrolase (OPH) activity (*k*_cat_/*K*_M_ < 10^2^ M^-1^s^-1^), which is similar to other previously studied DHCH enzymes^27^. In contrast, MPH-wt exhibits substantial activity against all four OP substrates (*k*cat/*K*M = 10^2^-10^4^ M^-1^s^-1^) and only moderate DHCH activity (*k*_cat_/*K*_M_ = 1.4 × 10^3^ M^-1^s^-1^). Interestingly, AncDHCH1 and AncDHCH2 exhibit higher OPH activity compared to the three extant DHCHs and AncDHCH3, albeit still 100-1000-fold lower than MPH-wt for three of the OP substrates (with ethyl-paraoxon being the exception) (**Fig. 1d** and **Supplementary Table 2**). These results suggest that promiscuous OPH activity in AncDHCH1 was a non-physiological and serendipitous promiscuous activity that arose prior to the introduction of methyl-parathion into the environment, and this promiscuous activity was recruited and then optimized in response to the appearance of this novel substrate.

### Five substitutions were responsible for the evolution of high methyl-parathion hydrolase activity

We sought to determine the minimal set of genetic changes that was responsible for the evolution of OPH activity. Out of a total of 32 changes (31 substitutions and one insertion) that occurred between AncDHCH1 and MPH-wt, we prioritized testing a subset based on two criteria: 1) the mutation is located in the vicinity of the active site, and 2) the residue otherwise exhibits high conservation either within the extant MPH orthologs or the DHCH orthologs (**Fig. 2**, and **Supplementary Fig. 4**). We identified five genetic changes that satisfied these criteria: four substitutions, *l*72R, *h*258L, *i*271T, and *f*273L (note that the *small italic letter* denotes the ancestral state for each amino acid residue while the large letter denotes the derived MPH state) and one insertion, Δ193S (**Fig. 2a-b**). We assessed the functional effect of these five mutations together by mutating each position in the AncDHCH1 genotype (AncDHCH1+m5) and also by reversing the five positions to their ancestral states in MPH-wt (MPH-m5). We then functionally characterized each genotype by measuring the purified enzyme’s activity against DHC and the four OP substrates. MPH-m5’s activity profile is almost identical to AncDHCH1, while AncCHCH1+m5 largely recapitulates the MPH-wt profile (**Fig. 2c** and **Supplementary Table 2**). These five changes together result in a 900-fold improvement in methyl-parathion activity and an 800-fold reduction in DHCH activity, resulting in an overall 700,000-fold shift in relative MPH/DHCH activities. Taken together, this minimal set of five genetic changes was both necessary and sufficient to cause the functional transition from DHCH to OPH enzymes.

**Figure 2.**
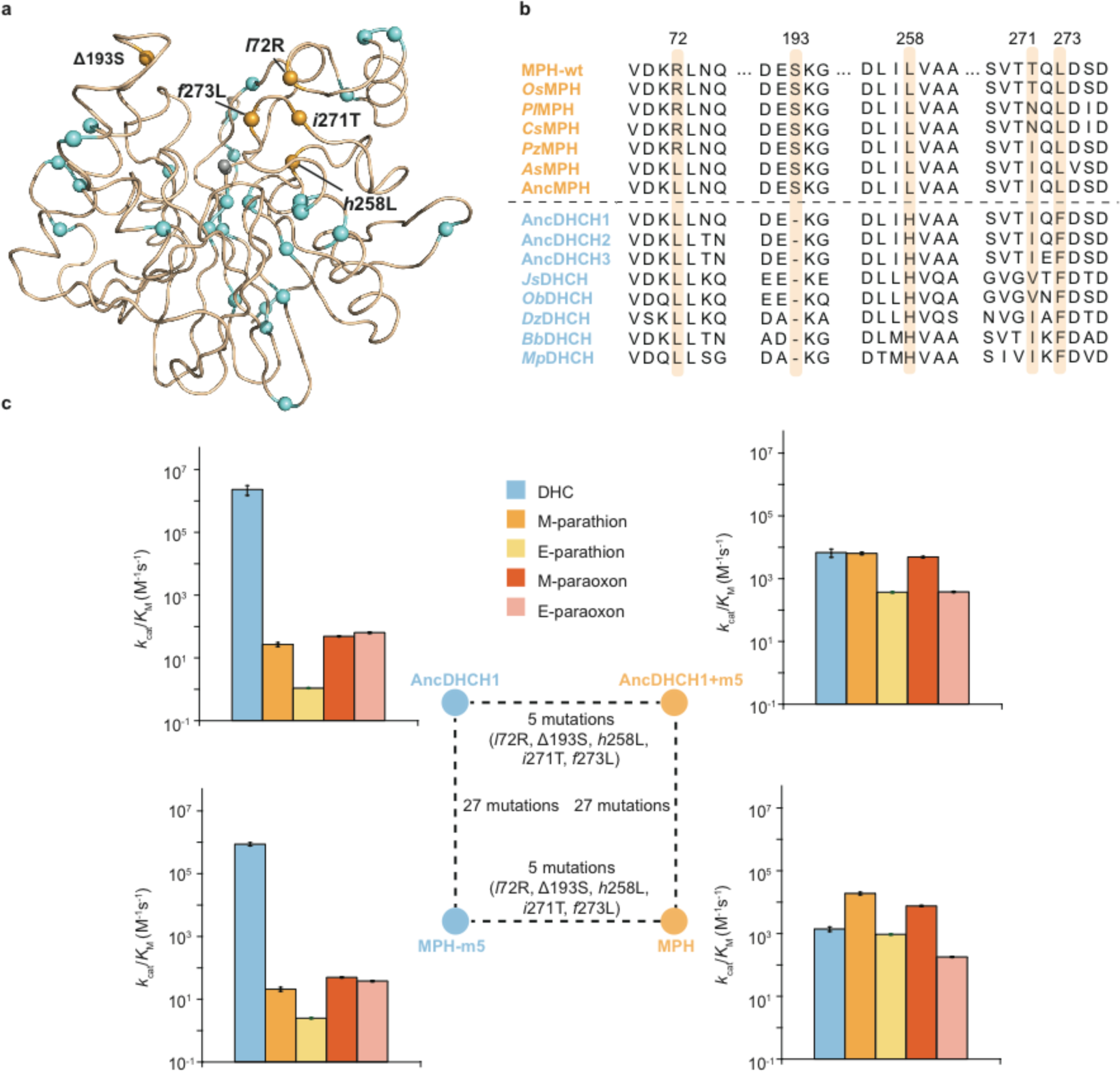
Identification of five key adaptive mutations between AncDHCH1 and MPH. **a**, Cartoon representation of the crystal structure of MPH-wt (PDB entry1P9E) with 32 mutations (spheres, 31 substitutions and 1 insertion) that occurred between AncDHCH1 and MPH-wt. Five active site mutations (*l*72R, Δ193S, *h*258L, *i*271T, and *f*273L) are highlighted as orange spheres. The two active site metal ions are shown as grey spheres. **b**, A cropped multiple sequence alignment of representative sequences of extant MPH, DHCH and resurrected ancestral enzymes. Residues at the positions where the five active site mutations have occurred between AncDHCH1 and MPH are highlighted in orange. A full multiple sequence alignment is presented in **Supplementary Fig. 4**. **c**, The effects of the five mutations on DHC and OP activities. Genetic relationships between the four enzymes are described in the inner square. A full description of the kinetic activities of the sequences can be found in **Supplementary Table 2**.

### MPH evolved specificity for methyl-parathion over other OP substrates

MPH’s evolved specificity is not only based on overall reaction chemistry, from C-O bond (DHCH) to P-O bond (OPH) cleavages, but also according to the substituents among the OP compounds, *e.g*., oxygen *vs*. sulphur (**Fig. 2c**). The ancestral enzymes (AncDHCH1 and MPH-m5) both exhibit an inverse linear relationship between log(*k*_cat_) and the partition-coefficient (log*P*) of the four OP substrates, with methyl substituents preferred over ethyl and oxygen over sulphur (**Fig 3a**). The five historical genetic changes, however, increase activity against sulphur-substituted substrates to a greater degree (∼1000-fold for methyl-parathion and ∼400-fold for ethyl-parathion) compared to oxygen-substituted substrates (∼200-fold for methyl-paraoxon and ∼10-fold for ethyl-paraoxon). Consequently, the log*P* dependence is lost in the derived enzymes (MPH and AncDHCH+m5), which no longer show preference for oxygen over sulphur, but still exhibit higher activities for methyl-substituents compared to ethyl (**Fig. 3a**). This indicates that the five mutations have specifically increased the enzyme’s ability to catalyze sulphur-substituted substrates.

To understand the structural basis for the functional changes, we solved the crystal structure of AncDHCH1 with a resolution of 1.7 Å (PDB entry: 6C2C) (**Fig. 3b** and **Supplementary Table 3**) and compared it to the previously published structure of MPH-wt (PDB entry: 1P9E). The two enzymes have almost identical overall structure, with the C-α backbone atoms aligned with a maximum r.m.s.d. of 0.46 Å, although there is a visible change in the shape of the active site pockets (**Figure 3b**). Previous work suggested that a terminal hydroxide ion bound by the α-metal ion in the active site carries out nucleophilic attack during OP hydrolysis^28^(**Supplementary Fig. 5**). The positions of the main catalytic machinery, including the active site metal ions and the residues that coordinate them, are nearly identical between AncDHCH1 and MPH (**Fig. 3b** and **Supplementary Fig. 6**). Four of the key mutations – *h*258L, *i*72R, *i*271L and *f*273L – are clustered in relatively close proximity to each other, and appear to function collectively to enlarge a potential binding pocket in the derived active site. The remaining mutation, Δ193S, is located on a loop opposite of the other four residues, and appears to have altered the conformation of the loop (**Figure 3b**). Docking of the methyl-parathion substrate into MPH indicates that the substrate is well accommodated in the more open active site of the derived enzyme, but encounters steric clashes with Phe273 in the ancestral state (**Fig 3c**). Taken together, the five mutations have enabled MPH to develop a better complementarity between the active site and the new substrate, methyl-parathion: MPH maintains a preference for methyl substituents over ethyl, but has evolved to become less discriminative between sulphur and oxygen substituents, whereas the interactions between the ancestral enzyme and OP substrates were governed to a greater degree by hydrophobic effects.

**Figure 3.**
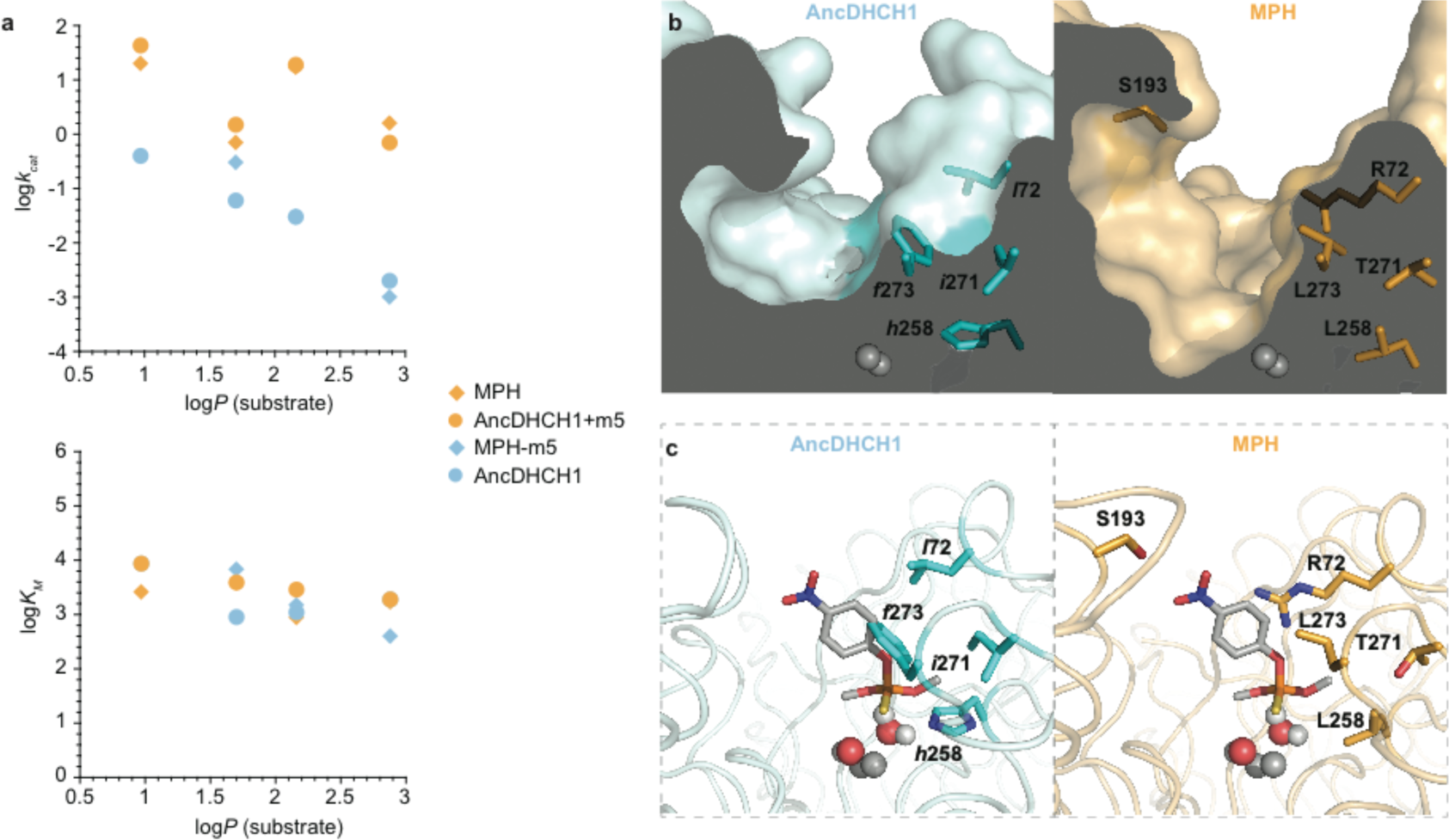
Structural and biochemical effects of five key mutations. **a**, A plot of Hansch hydrophobic constant (log*P*) versus *k*_cat_ (top) and *K*_M_ (bottom). Plots were generated using the activity profiles of the enzymes towards four different OP substrates: methyl-paraoxon (log*P* = 0.97), ethyl-paraoxon (log*P* = 1.7), methyl-parathion (log*P* = 2.16), and ethyl-parathion (log*P* = 2.88). Pale cyan indicates enzymes where the five key residues are in their ancestral states; light orange are enzymes with the derived state residues. **b**, Surface representation of the active sites of AncDHCH1 (PDB entry 6C2C) and MPH (PDB entry 1P9E) showing the change in cavity. The five key active site mutations are highlighted as sticks. The active site metals are shown as grey spheres. **c**, Molecular docking of the methyl-parathion substrate in the active sites of AncDHCH1 (left) and MPH (right). The active site metals are shown as grey spheres. The bridging and terminal (which acts as the likely nucleophile) water/hydroxide molecules are shown as spheres.

### A rugged adaptive landscape suggests a highly constrained trajectory during MPH evolution

We next sought to understand the evolutionary process that generated MPH activity. We characterized the topology of the adaptive landscape that encompasses the functional transition between AncDHCH1 and MPH by generating and assaying a complete combinatorial set of all five genetic changes (a total of 32 genotypic combinations) between MPH-m5 and MPH-wt (**Fig. 4a** and **Supplementary Table 5**). All variants were overexpressed in *E. coli* and the catalytic activity of the cell lysate for methyl-parathion was measured. Note that the soluble expression of all the genotypes is very similar, indicating that changes in the cell lysate activity largely reflect changes in catalytic efficiency and not the level of protein stability and expression (**Supplementary Fig. 7**). MPH-wt is found to be the single global maximum across this landscape, without any alternative local maxima. The five mutations, when introduced into MPH-m5, synergistically improve MPH activity by 970-fold, which exceeds the sum of the 210-fold improvement predicted from a null-additive model (*i.e.*, no epistasis) based on singular effect of each mutation in the background of the ancestral MPH-m5 (**Fig. 4a**). However, the adaptive landscape is rugged and only a limited set of adaptive trajectories is accessible: only 19 out of the 120 (5!) possible trajectories that connect MPH-m5 to MPH-wt could be completed without intermediate steps that required a reduction in MPH activity, indicating that the order in which those mutations accumulated was highly constrained and rather deterministic (**Fig. 4a**). This is because epistasis is prevalent, and all of the mutations display a broad range of effects depending on the presence or absence of other mutations (up to ∼600-fold variation in the effect of a single mutation depending on the genetic background in which it is introduced, **Fig. 4b**): three mutations (*l*72R, *i*271T, and *f*273L) exhibit sign epistasis, *i.e.*, a reversal in the direction of their effect, from being initially deleterious to eventually beneficial, while the remaining two mutations (*h*258L and Δ193S) exhibit strong magnitude epistasis, ranging from highly positive initially to almost neutral if introduced late in the trajectory. Notably, *i*271T is almost always deleterious, including when it is introduced into MPH-m5 (∼5-fold decrease), and becomes positive only after the prior appearance of three mutations, *l*72R/*h*258L/*f*273L (**Fig. 4b**). Given the environment in which MPH evolved (*i.e.*, with high amounts of methyl-parathion), we assume that, for a given genotype, the mutation that causes the greatest increase in OPH activity would be the most likely to be fixed. Following this assumption, we infer that the most likely order in which the five historical adaptive mutations were accumulated was *h*258L, Δ193S, *f*273L, *l*72R, and then *i*271T. Our proposed trajectory is also supported by the polymorphism observed in extant MPH genes at sites 72 and 271, where a few of the putative MPH orthologs still contain the ancestral residues, suggesting that they were most likely accrued at a later stage along the trajectory to the extant MPH-wt (**Supplementary Fig. 4**).

**Figure 4.**
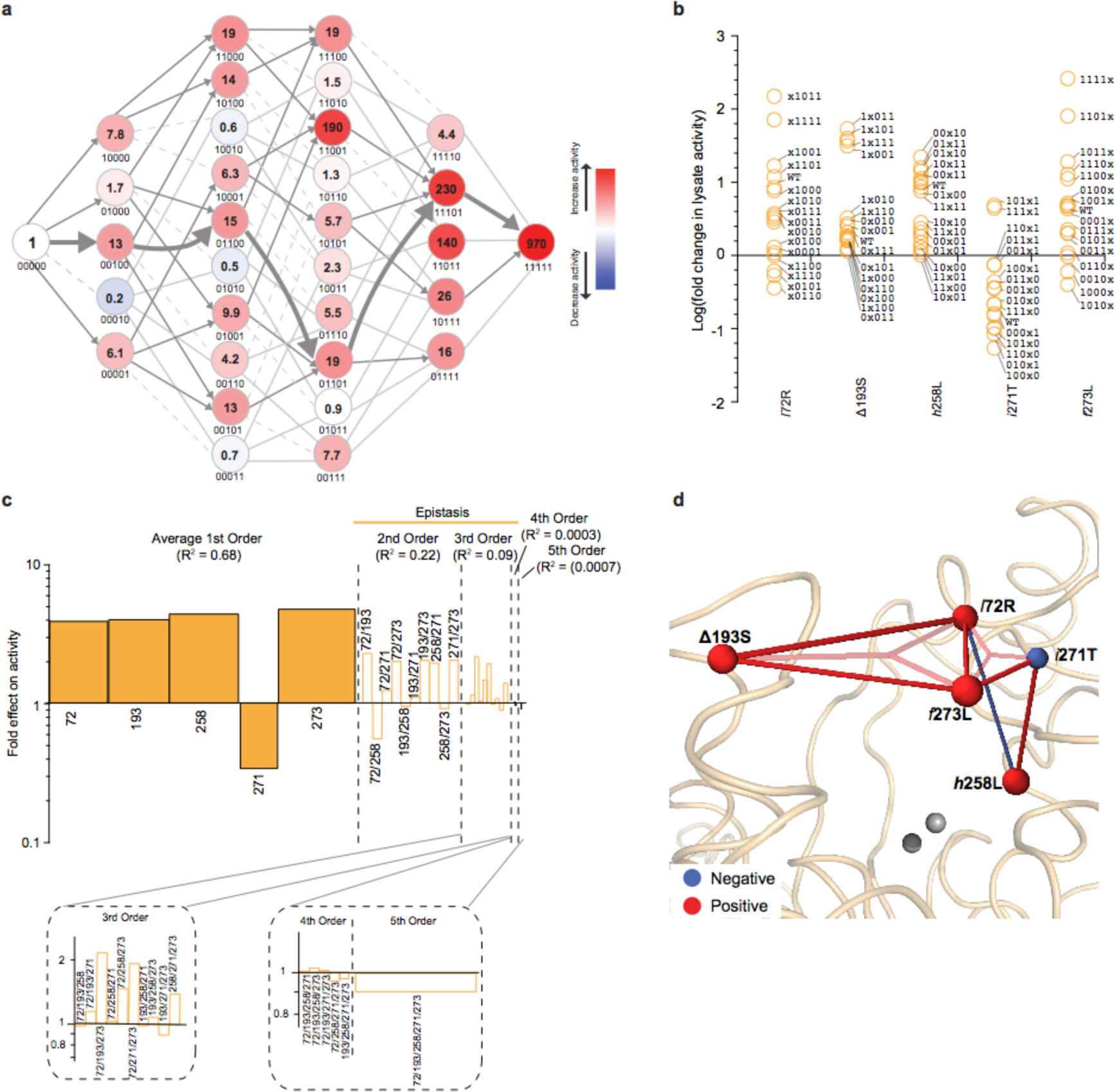
Adaptive landscape and mutational effects of key mutations for methyl-parathion hydrolase activity. **a**, The adaptive landscape between MPH-m5 and MPH-wt for methyl-parathion hydrolase activity. Each node represents a unique variant, with the genetic background indicated according to the numerical order of the residues (i.e., 72, 193, 258, 271, 273), where “0” refer to the ancestral and “1” refer to the derived state (e.g., 10000 denotes m5+*l*72R). Number in the centre of each node indicates its cell lysate activity relative to MPH-m5. Edges connecting nodes represent single mutational steps, with solid grey lines indicating paths that lead to an increase in fitness from the previous node and are evolutionarily accessible, dashed light lines paths that are inaccessible due to a decrease in fitness from the previous node, and solid grey lines paths that lead to an increase in fitness, but are inaccessible due to a decrease in fitness observed in a previous node. A full description of the activities of all the variants can be found in **Supplementary Table 5**. **b**, The singular mutational effects of each of the 5 mutations in 16 different genetic backgrounds. “x” indicates the mutation whose singular effects are plotted. **c**, Statistical analysis of mutational effects in the adaptive landscape. Mutations are indicated by their residue numbers (i.e., 72 denotes the effect of *l*72R; 72/193 denotes the effect of m5+*l*72R/Δ193S). The heights of the bars of the 1st order effects indicate the average effects (fold-change in lysate activity) of the five mutations. Widths of the individual bars in the 1st order effects correspond to the portion of variation in activity (R^2^ in the linear regression model) attributable to each of the singular effects. The heights of the bars in the 2nd, 3rd, 4th, and 5th order effects correspond to the average epistatic effects of the combined mutations. Widths of the 1st, 2nd, 3rd, 4th, and 5th order effects correspond to the variation attributable to combined mutational effects, calculated as the increase in adjusted R^2^ of the fit to the data when each term is added to a linear regression model. **d**, Graphical representation of the effect of the five mutations and their epistatic relationships mapped on the structure of the MPH (PDB entry 1P9E). Positions of the each residue reflect the configuration of the alpha-carbon of the residues in the MPH crystal structure. The size and colour of the nodes represent the magnitude of the average singular effects on MPH activity of the mutation. Edges connecting nodes indicate significant epistatic interactions between the residues [log_10_(epistatic effect) > 1]; solid lines indicate interactions between pairs while transparent lines indicate interactions between three residues.

### High-order epistasis shapes the adaptive landscape of MPH activity

To understand the molecular basis for the extensive epistasis and rugged adaptive landscape, we performed statistical analyses on the mutational effects by generating a series of nested linear models and fitting the landscape (**Fig. 4c)**. First, a simple non-epistatic model that determines the average “main effects” (1^st^ order effects) of each mutation across all genetic backgrounds (solid bars). We next constructed a series of more complex models that include pair-wise interaction effects (2^nd^ order), and then 3^rd^ order, 4^th^ order or 5^th^ order effects (see **Methods**). This linear modelling approach allows us to determine the “contribution” of each order of epistasis in terms of the improved R^2^ of each linear model’s fit to the experimental data^19,20^. Overall, epistasis accounts for more than 30% of the total variation in MPH activity across this evolutionary landscape. Interestingly, about 10% of the total variation is explained by high-order epistasis (3^rd^-order or higher) (**Fig 4c**). To verify the significance of this higher-order epistasis on the most likely sequence of mutations, we reconstructed the adaptive landscape using the parameters obtained in the linear models that include only lower-order effects (**Supplementary Fig. 8** and **Supplementary Table 6**). When only the 1^st^ and 2^nd^ order effects are included, the reconstructed landscape differs significantly from the experimental data. By contrast, when all the epistatic effects up to the 5^th^ order are included, the reconstructed landscape roughly matches the experimental data. This further bolsters the importance of high-order epistasis in determining the pathways that are available.

To further decipher the molecular basis for epistasis, we visualized all 2^nd^-and 3^rd^-order epistatic interactions above a minimum threshold [log_10_(epistasis effect)>1] as “edges” between the Cα for each of residues involved on the structure of MPH (**Fig. 4d**). This analysis unveiled a highly connected interaction network among the five mutations. 2^nd^ order epistasis was observed between *l*72R and *f*273L, as well as *h*258L and *i*271T. In several cases, the pairs of residues are in relatively close proximity to each other in the active site, and may form direct physical interactions. In other cases, however, epistasis is also observed between residues that are physically far apart: for example, *h*258L exhibits antagonistic epistasis with *l*72R in spite of being located almost 7 Å away (**Figure 4d** and **Supplementary Fig. 9**). Even more interestingly, Δ193S exhibits synergistic epistasis with *l*72R and *f*273L via both 2^nd^-and 3^rd^-order epistasis despite being located on the opposite side of the active site (>10 Å distance). We hypothesize that these long-range epistatic interactions are likely mediated by the substrate itself, and that the mutations improve catalytic activities by altering the substrate position in the active site.

### Mutational epistatic effects change dramatically between O-vs. S-substituted substrates

We further conducted comparative landscape analyses between the four different OP substrates to determine how MPH evolved such high substrate specificity. We assayed all 32 genotypic variants against the remaining three OP substrates (ethyl-parathion, methyl-paraoxon, and ethyl-paraoxon) and quantified the epistatic interactions using the same linear modelling approach described above (**Fig. 5**, and **Supplementary Fig. 10**). The topology of the adaptive landscapes and the main and epistatic effects of the mutations for methyl-*vs.* ethyl-parathion are largely similar (**Fig 5a** and **Supplementary Fig. 10a**); consequently, the structural epistasis interaction networks for these two substrates are virtually identical (**Fig. 5d**). Comparing the effects of each mutation across all 16 possible genetic backgrounds between the two substrates reveals a strong linear correlation (R^2^ = 0.96), again indicating that epistatic interactions are similar in the two backgrounds; however, three mutations, *l*72R, Δ193S and *f*273L, exhibit a slight preference for the methyl substituent over ethyl (by roughly 2-fold) (**Fig. 5a** and **Supplementary Fig. 11a**).

**Figure 5.**
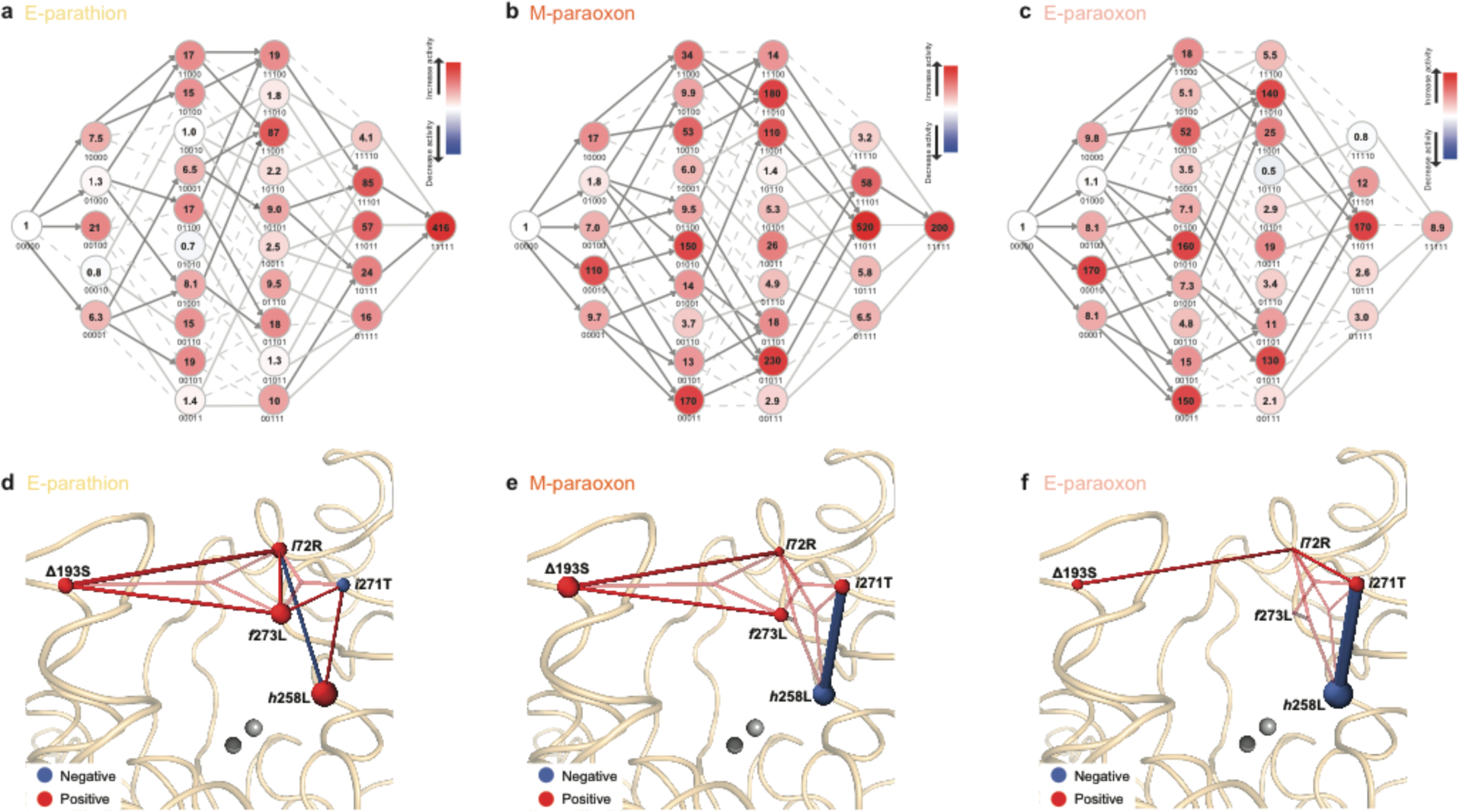
Adaptive landscapes and mutational effects for three other OP substrates. **a-c**, The adaptive landscape of 32 variants between MPH-m5 and MPH-wt for **a**, ethyl-parathion, **b**, methyl-paraoxon, and **c**, ethyl-paraoxon. Each node represents a unique variant with the number at the centre representing its cell lysate activity relative to MPH-m5. Edges connecting nodes represent single mutational steps. The definition of the colour of the edges is the same scheme as in **Figure 4**. A full description of the activities of all the variants can be found in **Supplementary Table 5**. **d-f**, A graphical representation of the effect of the five mutations and their epistatic relationships mapped on the structure of the MPH (PDB entry 1P9E). Positions of each residue reflect the configuration of alpha-carbon of the residues in the MPH crystal structure. The colour scheme for the nodes and edges are described in **Figure 4**. A full description of the statistical characterization is described in **Supplementary Fig. 10**.

By contrast, the discrimination between sulphur *vs.* oxygen substituents revealed dramatic differences in the adaptive landscape for activity against otherwise identical substrates. The landscapes for paraoxons are noticeably more rugged, containing several local maxima (**Fig. 5b-c**). In particular, the derived MPH-wt genotype is not the most active genotypic variant for paraoxons; instead, MPH-m5+271 and MPH-m5+72/193/271/273 are the two most active variants against ethyl-paraoxon (**Fig. 5c**). The main and epistatic effects of each mutation for the two paraoxon substrates differ significantly compared to that of the parathion substrates (**Supplementary Fig. 10b**-**c**); consequently, the interaction networks for the paraoxons are substantially different from that of parathions, with the paraoxon networks exhibiting fewer significant 2^nd^-order and more 3^rd^-order epistatic interactions (**Fig. 5e-f**). Most notably, the average effect of *h*258L has switched from being positive in parathion to negative in paraoxon; conversely the average effect of *i*271T has changed from negative in parathion to positive in paraoxon. Moreover, the 2^nd^ order effect of *h*258L-*i*271T has changed from being synergistic in the parathion backgrounds to strongly antagonistic in the paraoxon backgrounds (**Fig. 5e-f** and **Supplementary Fig. 10b-c**).

### Adaptive landscapes uncover the molecular mechanisms that underlie OP substrate specificity

Paraoxons and parathions are identical in overall size and structure; thus, the observed differences in enzyme specificity must stem from the phosphoryl sulfur or oxygen substituent. The reduced electronegativity of sulphur results in a less polarized, more hydrophobic molecule. Although this results in slower base-catalyzed hydrolysis in solution^29^, the chemical reactivity of the oxons and thions with highly reactive 4-nitrophenol leaving groups is not substantially different in low-dielectric environments such as gas-phase simulations or enzyme active sites^30^. Moreover, the *k*_cat_ between substrates with methyl and ethyl sidechains also differ, despite the fact that this will not affect the reactivity of the molecules (**Supplementary Table 2**).

In our aforementioned molecular docking experiment, the wild-type dihydrocoumarin hydrolase has poor complementarity to the phosphotriesters; the mutations appear to have made the active site substantially more accommodating to the new substrates (**Fig. 3c**). In particular, *h*258L appears to increase the size of the binding pocket in order to accommodate the methyl/ethyl-side chain of the substrate and *i*271T, which is located adjacent to *h*258L, may affect the position of the His/Leu258 through steric hindrance of some rotamers. A comparison of the effects of each mutation between parathions and paraoxons reveals that the effects of *h*258L and *i*271T are strongly dependent on the state of the other residue (**Fig. 6a** and **Supplementary Fig. 11b-c**): the average effect of *h*258L is mildly positive for both parathion and paraoxon when position 271 in the ancestral state (*Ile*). However, it becomes more positive for parathion and strongly negative for paraoxon when position 271 is in the derived state (Thr) (**Fig. 6c**). Conversely, the average effect of *i*271T is deleterious for parathion and positive for paraoxon when the position 258 is in the ancestral state (*His*), but becomes deleterious for both substrates when 258 is in the derived state (Leu) (**Fig. 6c**). Interestingly, a plot of the double mutational effects of *h*258L-*i*271T between methyl-parathion and methyl-paraoxon in the eight genetic backgrounds exhibits a linear correlation (**Fig 6b**, R^2^= 0.93), indicating that there is variation in the strong epistasis between *h*258L and *i*271T that depends on the ligand. The other three mutations also exhibit a greater positive effect for parathion over paraoxon (**Fig 6a** and **Supplementary Fig. 11**), indicating that all mutations collectively contribute to the preference for sulphur, resulting in the derived MPH having a >5-fold higher improvement in methyl-parathion activity over methyl-paraoxon. Taken together, our results suggest that the improved activity for sulphur-substituted substrates in the derived enzyme emerges gradually due to the collective effect of all five mutations, which must arise in the right order because of the complex interaction network that governs molecular recognition across this evolutionary transition.

**Figure 6.**
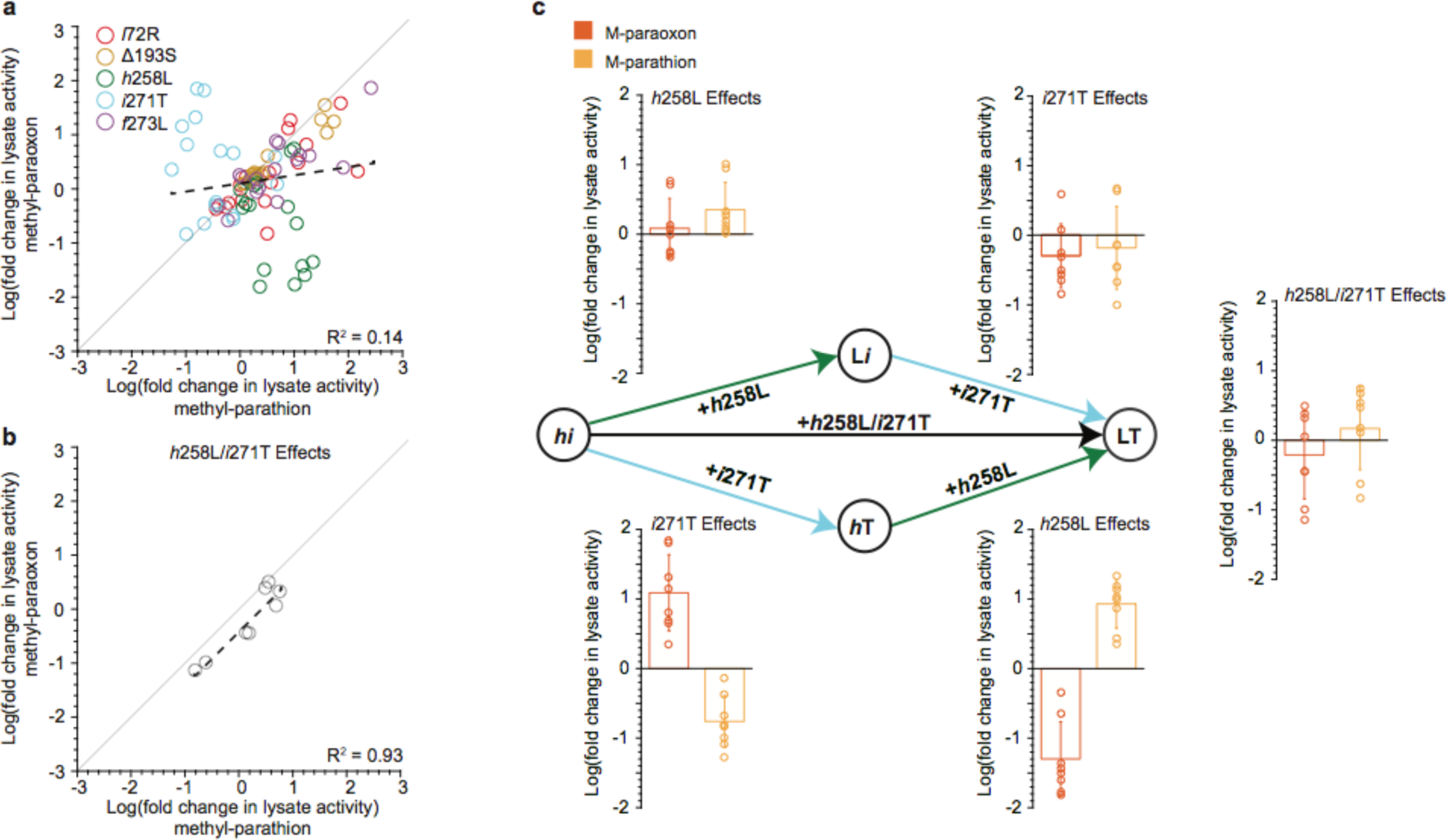
Changes in the singular and epistatic effects of mutations between methyl-parathion and methyl-paraoxon substrates. **a** and **b** show the effects of **a**, each of the five key mutations in all 16 possible genetic backgrounds and **b**, The combined effects of *h*258L-*i*271T in all 8 genetic backgrounds for methyl-paraoxon against that of methyl-parathion. The dashed line indicates the linear fit, with the R^2^ indicated in the bottom right corner of each plot. The solid black line running through the graph represents a slope of 1. **c**, The pairwise epistatic effects of *h*258L and *i*271T on methyl-parathion and methyl-paraoxon substrates. Labels in the centre of the nodes indicate the genotypes at the two positions, with the mutation occurring between each genotype indicated on the arrow. The singular effects of *h*258L and *i*271T in the 8 possible genetic backgrounds where one of the residues is in the ancestral state (going from the *hi* genotype) or in the derived state (going from either the L*i* or *h*T genotype), along with the pairwise effects of *h*258L-*i*271T, are plotted for methyl-paraoxon (red) and methyl-parathion (yellow). The dots on the plots represent the effects of the mutation(s) across the different genetic backgrounds, the heights of the bars represent the average of these effects, and the whiskers indicate the standard deviation.

Interestingly, the improved complementarity of the enzyme for the substrates has little effect on the binding affinity: the worse substrates (thions) actually have a lower *K*_M_ value (better formation of Michaelis complex) than the oxons, and the *K*_M_ values change little over the course of the evolution. In fact, almost all of the change in activity can be attributed to improvements in *k*_cat_, with a more beneficial effect observed for the thion substrates than the oxon substrates. To investigate the cause of the differential improvement in thion over oxon substrates, we analysed how they might affect the enzyme mechanism (**Figure 3**). None of the five mutations are in a position to participate any direct interactions with the terminal oxygen/sulphur, which loosely associates with the active site metal ions, nor do they directly affect the catalytic machinery (nucleophile generation or leaving group departure) (**Fig 3c**). Thus, the changes in activity don’t appear to be from changes in the chemistry of the reaction, but are more likely the result of changes in the nature, or productivity, of the substrate binding. In the ancestor, where there is poor steric complementarity between enzyme and substrate, the more polar oxons are turned over at ∼10-fold greater rates than the more hydrophobic thions. In contrast, in the evolved enzymes, which do have substrate-shape complementarity to align the substrates for hydrolysis, the turnover rates are comparable, as would be predicted from their intrinsic reactivity in low-dielectric environments. This suggests that while overall substrate binding (as inferred through inspection of *K*_M_) does not change markedly through evolution, the quality (or productivity) of substrate binding, i.e. whether the substrate is bound in a conformation aligned for hydrolysis, has improved.

## Discussion

Previous studies using ASR examined functions that evolved millions or billions of years ago^2–4,31–35^. Our study demonstrates that this technique, combined with biochemical and mutational assays, can effectively uncover the molecular mechanisms underlying recently evolved functional novelty. The introduction of xenobiotics into the environment led to the evolution of many novel enzymatic functions^36–38^. While numerous xenobiotic-degrading enzymes have been functionally characterized, the evolutionary origins and dynamics for most of these new sequences are still unknown due to large genetic “gaps” that currently exist in the databases^38–40^. Metagenomic sequencing projects may continue to fill in these gaps by uncovering enzyme sequences that are more ancestral-like^41^. As more and more novel sequences are obtained, ASR is becoming an increasingly valuable tool for researching functional evolution.

Our observations of the mechanisms underlying the evolution of MPH are consistent with the conclusions of several previous studies on protein evolution. Specifically, efficient MPH enzymes emerged rapidly, via a handful of genetic changes, by optimizing a promiscuous activity present in its ancestral state^3,4,27,35,42,43^. At the same time, epistasis between key adaptive mutations is prevalent, and acts to constrain the evolutionary pathways that were available^10,14,15,17,18,21,22^; early mutation(s) played a permissive role by epistatically generating or enhancing the positive effect of later mutations. As a consequence, the sequence in which historical substitutions accumulated likely occurred in a deterministic fashion^44–47^.

Most previous studies characterizing adaptive fitness landscapes have focused on unveiling the mutational epistasis that constrain the accessibility of evolutionary pathways^10,14,15,17,18,22^. Our in-depth statistical analyses of the adaptive landscapes of five key mutations, combined with robust biochemical and biophysical characterization, provided deep insight into the molecular mechanisms underlying the optimization of methyl-parathion activity. Moreover, we developed a novel approach in which we compare the adaptive landscapes for multiple substrates to decipher the interactions between mutations and substrate substituents that enabled MPH to increase recognition specifically for sulphur-substituted substrates. Our results suggest that the MPH active site improved complementarity to methyl-parathion through the collective effects of five mutations that form a complex and interconnected network. We speculate that each mutation impacts catalysis by reshaping the active site to better accommodate the new substrate and by reducing non-productive modes of binding, while also reorienting the substrate to optimize the nucleophilic attack and transition state stabilization. It is probable that higher turnover rate of the oxons in the non-complementary ancestor are a consequence of the more polarized P=O bond resulting in an increased likelihood of productive coordination to the active site metal in the absence of a complementary binding site, in comparison to the more hydrophobic thions that would be more likely to coordinate non-specifically through van der Waals interactions. The effects of non-productive binding have previously been described many proteins, including chymotrypsin and an unrelated bacterial phosphotriesterase^30,48^. This is also reminiscent of recent work that demonstrated how laboratory evolution of an arylsulfatase was able to cause a 100,000-fold increase in phosphonate-monoester hydrolase activity by enlarging the active site and repositioning the substrate^49^. In such cases, the structural consequences of mutations can be subtle in the active site, with sub-Ångström distance and angle adjustments resulting in optimized catalytic machinery, causing profound changes in catalytic efficiency and specificity. While the extent to which this model can account for the molecular mechanisms for novel enzyme functions more broadly remains to be seen, it is consistent with other models of enzyme dynamics and conformational diversity^50–53^. Fully assessing this will require a detailed understanding of the molecular relationships between the active site and the substrate in other examples of enzyme functional evolution. Combined with comprehensive and deep mutational analyses^13,16,20^, studies such as this can deepen our understanding of the molecular mechanisms that underlie the evolution of new protein functions.

## Acknowledgements

We thank A. Pabis for performing computational analysis and providing revision and comments on the manuscript. N.T. and E.B. thank the Human Frontier Science Program Research Grant. N.T. thanks the Natural Sciences and Engineering Research Council of Canada (NSERC) Discovery Grant (RGPIN 418262-12 and RGPIN 2017-04909). N.T. is a CIHR new investigator and a Michael Smith Foundation of Health Research (MSFHR) career investigator. S.C.L.K. thank the Knut and Alice Wallenberg Foundation, the Wenner-Gren Foundations and the Swedish National Infrastructure for Computing (SNIC). D.W.A thanks NSERC and the MSFHR for post-doctoral support.

## Author contributions

G.Y. and N.T. conceived and designed this study. G.Y. and F.B. performed activity assays and mutational analysis. D.A. performed statistical analyses. E.B. supervised bioinformatics. E.D. performed ancestral sequence reconstruction. F.B., N.H., and P.D.C. collected structural data. S.C.L.K. designed computational analysis. G.Y. and N.T. wrote the paper with input from all authors.

