## Supplementary information for "Higher-order epistatic networks underlie the evolutionary fitness landscape of a xenobiotic-degrading enzyme"

### Supplementary Figure 1

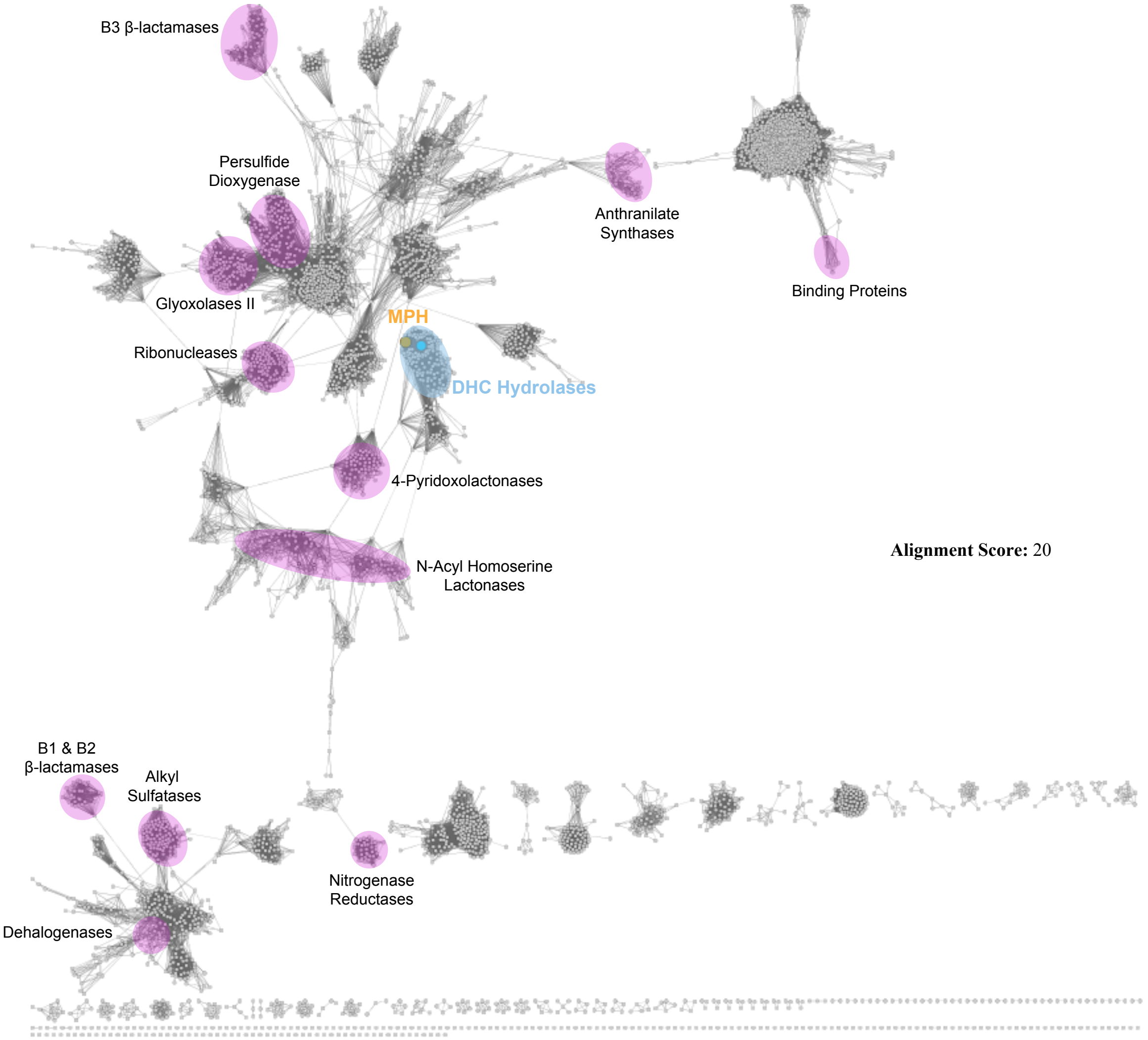


**Supplementary Figure 1. Representative sequence similarity network (SSN) of enzymes in the metallo-β-lactamase (MBL) superfamily generated using EFI-EST.** 37,709 unique protein sequences were obtained from the Uniprot database in March 2016, and the similarities among the sequences calculated using "all-vs.-all" BLAST pairwise comparisons. Sequences that share >40% sequence identity are grouped together as one node (circle) in the network, resulting in a total of 4,548 nodes representing all 37,709 sequences. Nodes are connected by an edge if the mean pairwise BLAST alignment score between all sequences in the node is above a threshold of 20. Sequence clusters that include experimentally characterized enzymes are highlighted in violet to indicate distinct functional families. The dihydrocoumarin hydrolase (DHCH) family is highlighted in cyan, with large coloured nodes in the family representing experimentally characterized sequences: the large cyan node represent characterized DHCH enzymes and the large orange node methyl-parathion hydrolases (MPH).

### Supplementary Figure 2

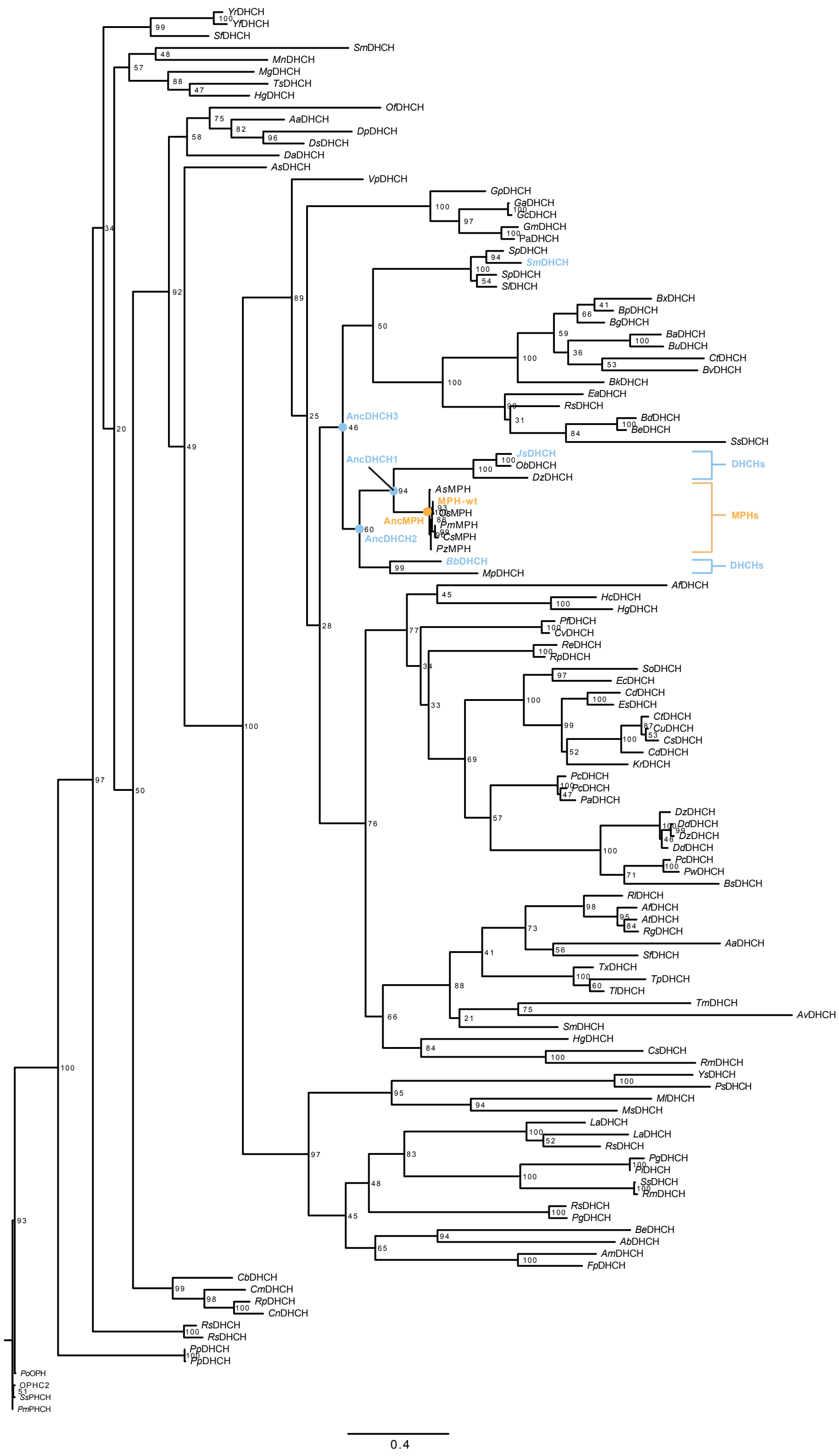


**Supplementary Figure 2. Phylogenetic tree of MPHs and representative DHCH enzymes.** A total of 153 of the closest MPH homologous sequences were used for the construction of the phylogeny via RAxML utilizing the LG protein model. Numbers at each of the branching nodes indicate bootstrap values. The cluster of MPH enzymes consist of a number of highly similar sequences (>90% sequence identity); the enzyme isolated from *Pseudomonas* sp WBC-3 (MPH-wt) is utilized for this study and labeled in orange. DHCH enzymes that were selected for characterization are labeled in cyan. Predicted ancestral sequences that were synthesized and characterized in this study are indicated by the cyan nodes. The most recent ancestral enzyme of the MPH (AncMPH) sequences is indicated by an orange node.

### Supplementary Figure 3

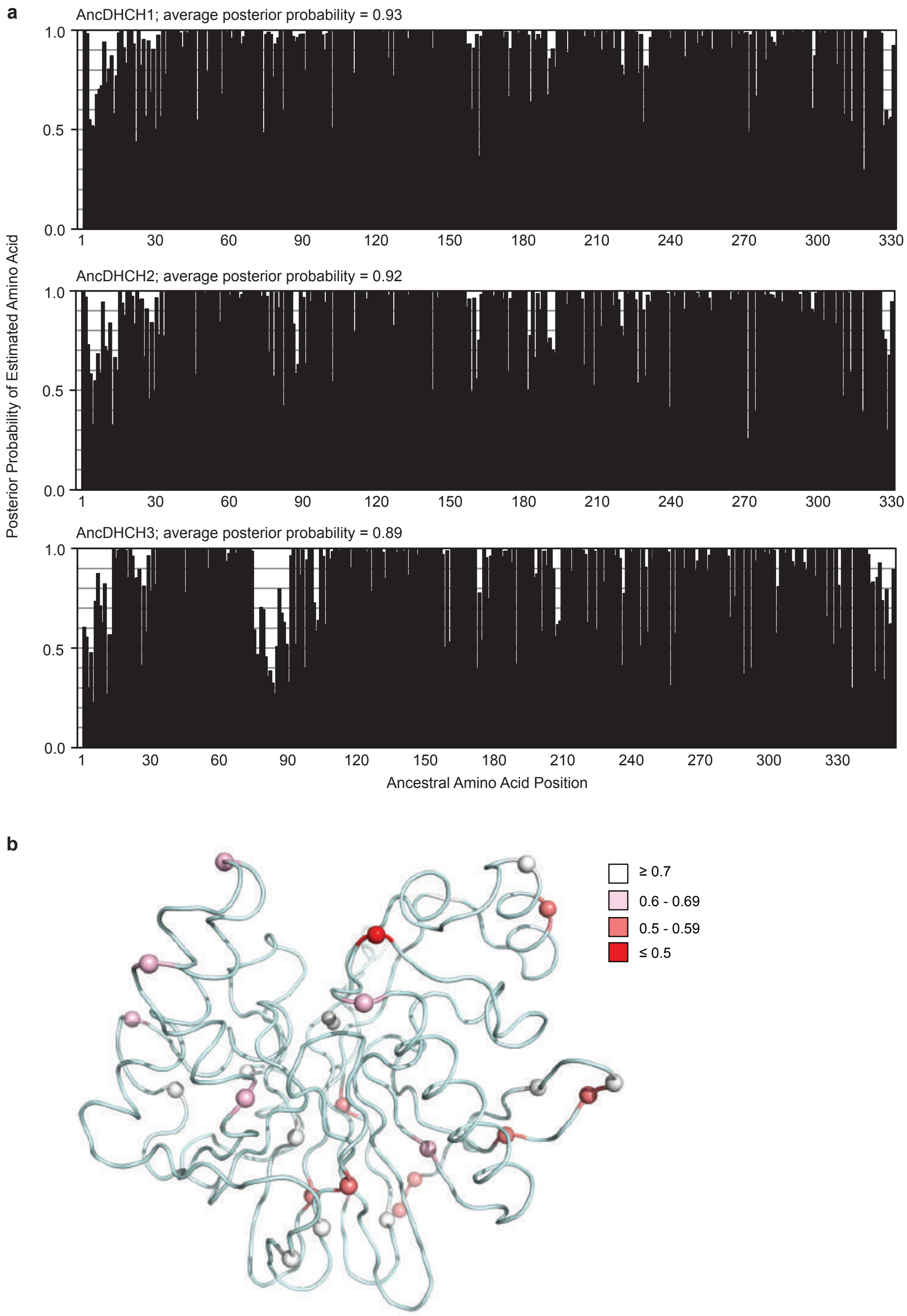


**Supplementary Figure 3. Site-specific posterior probabilities for ancestral amino acid sequence characterized in this study.** **a**, The Bayesian posterior probabilities of the most likely predicted ancestral amino acid at each position in the resurrected sequences of AncDHCH1 (top), AncDHCH2 (middle), and AncDHCH3 (bottom). The average posterior probabilities for each sequence is shown at the top of each graph. **b**, Cartoon representation of the structure of AncDHCH1 with the locations of ambiguous positions (predicted ancestral residues with a Bayesian posterior probability <0.8) are depicted as spheres. Colours of the spheres depict the posterior probability of the predicted residue, with red indicating lower probability and white higher probability. A full summary of the ambiguous positions and their posterior probabilities can be found in **Supplementary Table 1**. The two metal ions coordinated in the active site are shown as grey spheres.

### Supplementary Figure 4

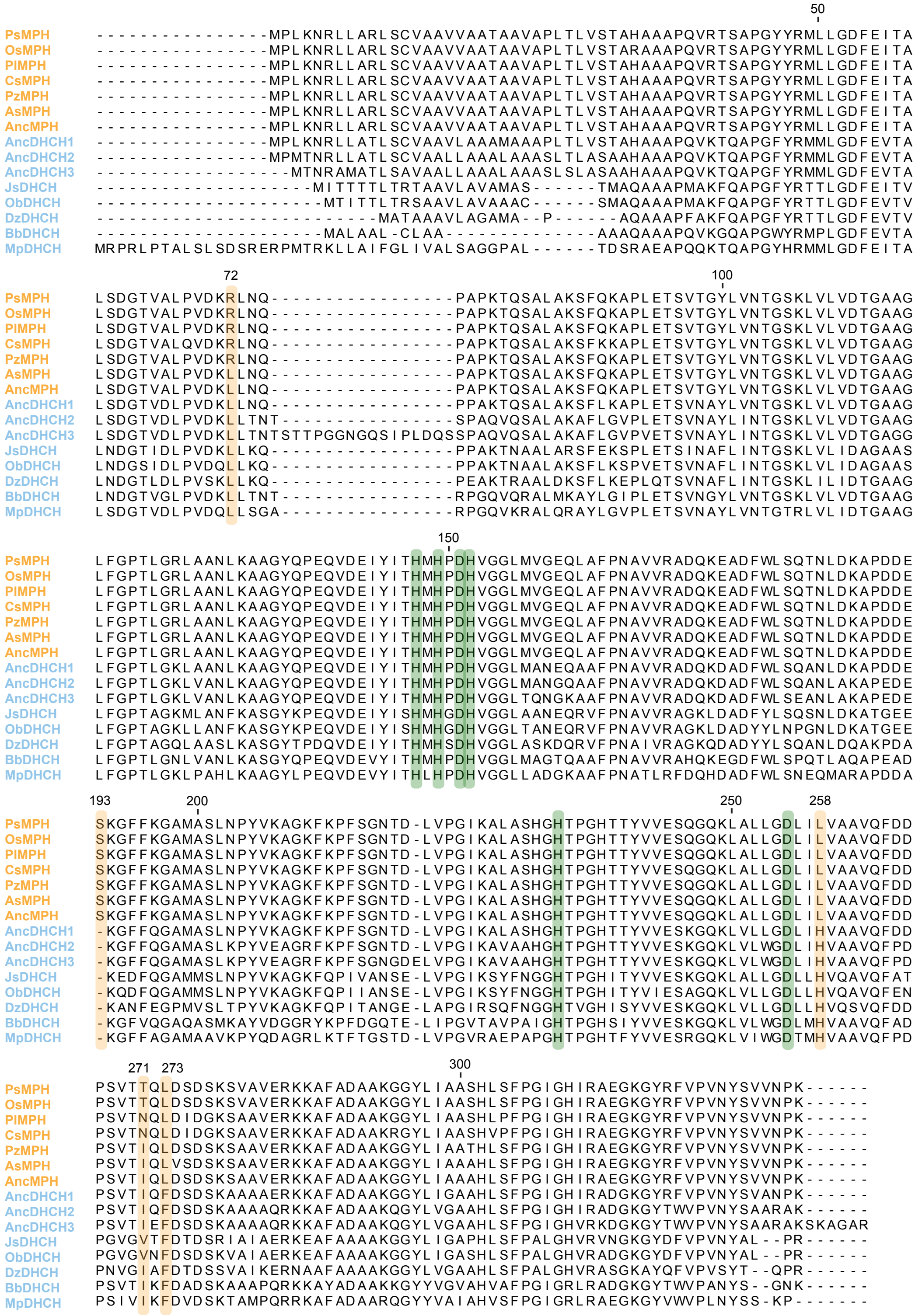


**Supplementary Figure 4. A multiple sequence alignment of representative extant MPH, DHCH, and predicted ancestral enzymes.** Residue numbering is based on the sequence of the structure of the MPH from *Pseudomonas* sp WBC-3 (MPH-wt, PDB ID: 1P9E). Extant MPH enzymes are labeled in light orange; extant DHCH enzymes and predicted ancestral sequences are labeled in pale cyan. Positions of metal binding residues are highlighted in green; positions of the five key functional mutations in MPH evolution are highlighted in orange.

**Supplementary Figure 5**

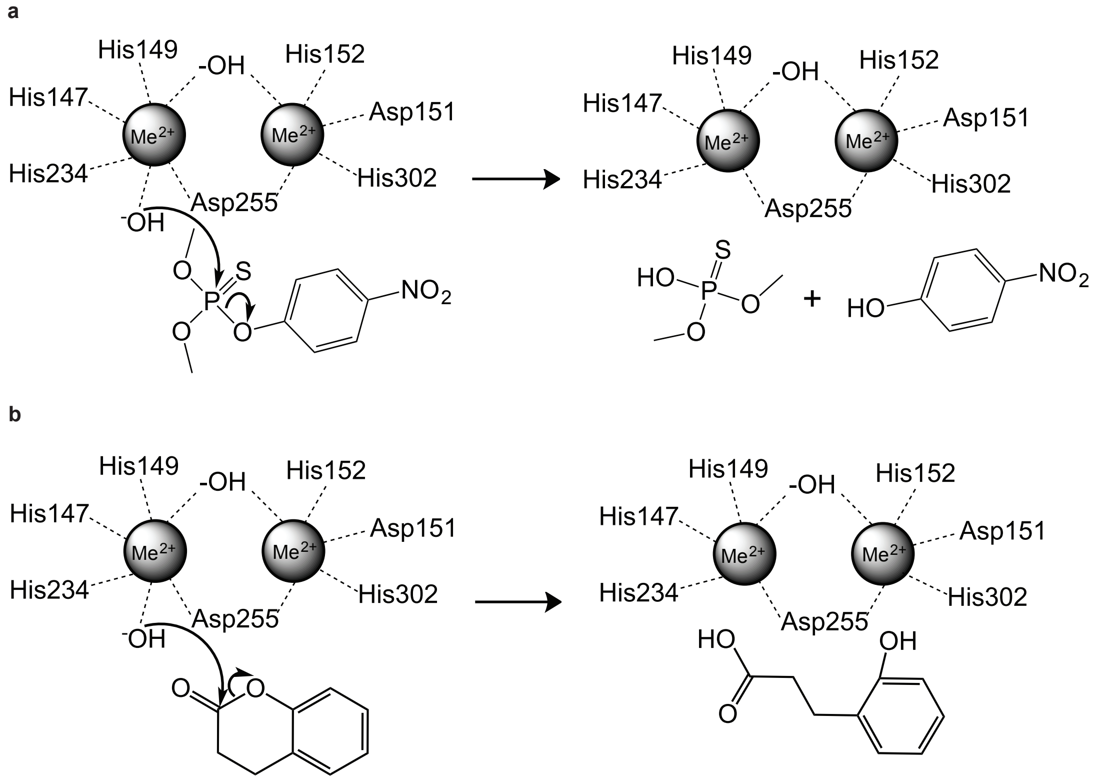


**Supplementary Figure 5. Proposed catalytic mechanisms of MPH for organophosphate and dihydrocoumarin hydrolytic reactions. a**, A proposed mechanism for OP hydrolysis of MPH as described by Purg *et al.*[28]. Metal cations (Me) in the active site are coordinated by adjacent histidines and aspartic acids, and a terminal hydroxide ion, which serves as a nucleophile that attacks the phosphate of the substrate. **b**, A proposed mechanism for DHC hydrolysis of MPH. The hydroxide ion that serves as the nucleophile in OP hydrolysis is presumed to play the same role in DHC hydrolysis.

**Supplementary Figure 6**

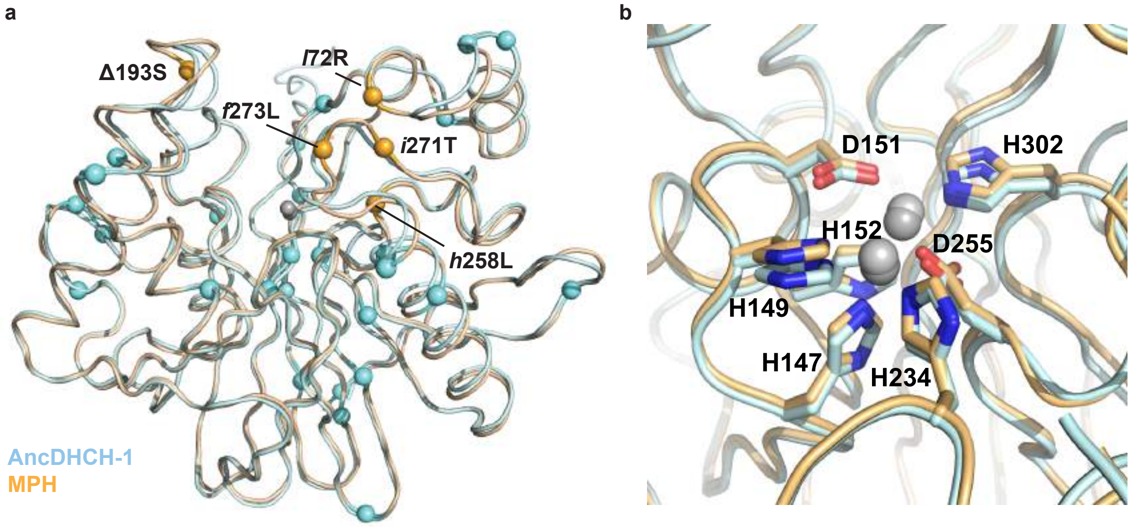


**Supplementary Figure 6. Comparison of the crystal structures of AncDHCH1 and MPH. a**, Overlay of the cartoon representations of the crystal structures of AncDHCH1 (pale cyan, PDB entry 6C2C) and MPH (light orange, PDB entry 1P9E). 27 of the 32 amino acid substitutions that have occurred between the two enzymes are depicted as cyan spheres. 5 amino acid mutations (*l*72R, Δ193S, *h*258L, *i*271T, and *f*273L) that occurred in the vicinity of the enzyme active sites are highlighted as orange spheres. The two metals coordinated in the enzyme active sites are shown as grey spheres. **b**, A close up view of the enzymes' active sites of the overlaid crystal structures of AncDHCH1 and MPH from **a**. Metal binding residues are highlighted as sticks. Metals bound in the active sites are shown as grey spheres.

**Supplementary Figure 7**

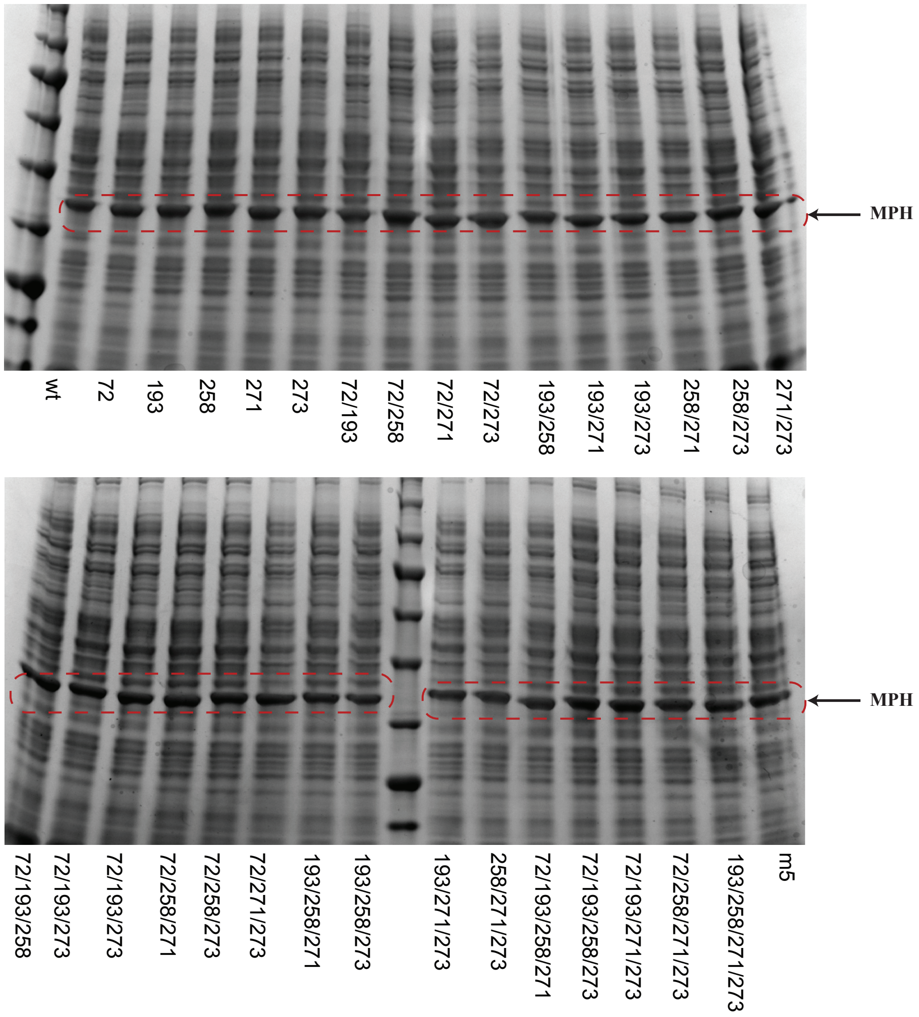


**Supplementary Figure 7. SDS-PAGE analysis showing soluble fractions of the 32 MPH combination variants.** The band corresponding to the Strep-tagged MPH enzyme (33.2 kDa) is indicated by dashed red circles. The genotype of each unique variant is indicated by numbers corresponding to the positions of the 5 key mutations on the background of the ancestral (m5) state (*i.e.*, 72 denotes m5+*l*72R mutation; 72/193 denotes m5+*l*72R/Δ193S).

### Supplementary Figure 8

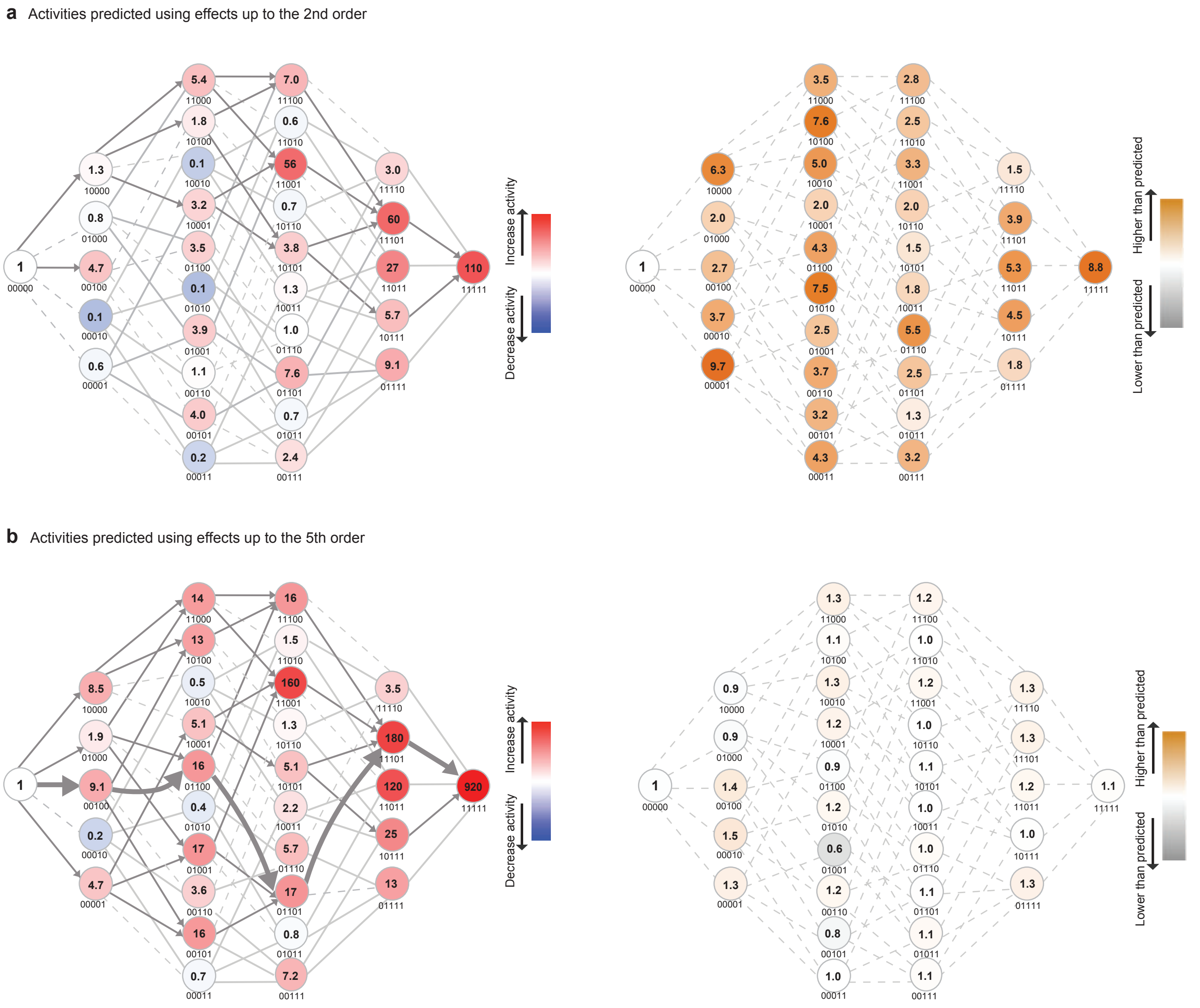


**Supplementary Figure 8. Reconstructed adaptive fitness landscapes of MPH for methyl-parathion activity using simulated data, where the activities of the 32 genotypes are predicted in linear regression models using a**, only 1st and 2nd order effects and **b**, using all epistatic effects up to the 5th order. The panels on the left show the fitness landscape for methyl-parathion. Each node represents a unique variant, with the genetic background indicated according to the numerical order of the residues (*i.e.*, 72, 193, 258, 271, 273), where “0” refer to the ancestral and “1” refer to the derived state (*e.g.*, 10000 denotes m5+*I*72R). Number in the centre of each node indicates its cell lysate activity relative to MPH-m5. Lines connect nodes that are separated by single mutations. Solid dark grey lines indicate mutational pathways that are evolutionarily accessible (results in an increase in fitness from the previous node). Dashed light grey lines indicate pathways that are inaccessible due to a decrease in fitness from a previous node. Solid grey lines indicate pathways that lead to an increase in fitness, but are evolutionarily inaccessible due to a decrease in fitness observed in the previous node. The panels on the right show the difference in activity between the simulated landscape and the actual, with the numbers on each node depicting the ratio between the actual and predicted fold change in activities. The dashed grey lines indicate nodes that are separated by single mutations. A complete summary of the simulated data is presented in **Supplementary Table 6**.

### Supplementary Figure 9

**a**

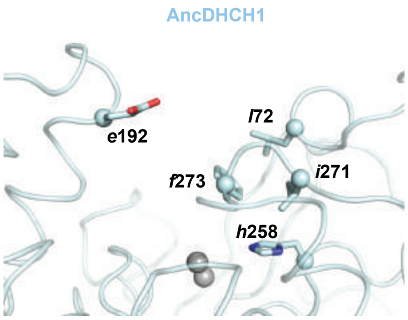


**AncDHCH1**

| | Distance between nearest atoms (Å) | | | | |
|---|---|---|---|---|---|
| | ***l*72** | ***e*192** | ***h*258** | ***i*271** | ***f*273** |
| ***l*72** | | 9.6 (C5-O) | 7.3 (C5-C5) | 4.1 (C3-C5) | **3.8 (C5-C4)** |
| ***e*192** | | | 13.4 (C2-Nτ) | 11.8 (O-C3') | 8.2 (O-C6) |
| ***h*258** | | | | **3.7 (Nπ-C4)** | **3.9 (C5-C2)** |
| ***i*271** | | | | | **3.9 (C5-C2)** |
| ***f*273** | | | | | |

| | Distance between Cα atoms (Å) | | | | |
|---|---|---|---|---|---|
| | ***l*72** | ***e*192** | ***h*258** | ***i*271** | ***f*273** |
| ***l*72** | | 14.4 | 10.1 | 5.6 | 7.6 |
| ***e*192** | | | 18.6 | 14.9 | 10.1 |
| ***h*258** | | | | 7.9 | 9.4 |
| ***i*271** | | | | | 5.4 |
| ***f*273** | | | | | |

**b**

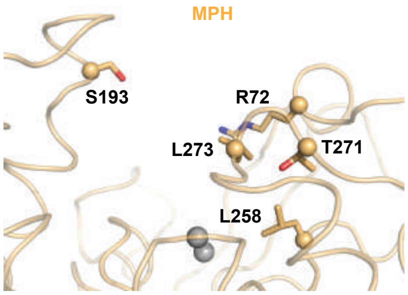


**MPH**

| | Distance between nearest atoms (Å) | | | | |
|---|---|---|---|---|---|
| | **R72** | **S193** | **L258** | **T271** | **L273** |
| **R72** | | 8.9 (Nδ-O) | 6.9 (Nδ-C5) | 5.3 (Cα-C4) | **3.6 (Nδ-C5)** |
| **S193** | | | 15.4 (O-C5) | 14.4 (O-O) | 9.1 (O-C5) |
| **L258** | | | | **3.6 (C5-O)** | 4.4 (C5-C5) |
| **T271** | | | | | **3.4 (O-C3)** |
| **L273** | | | | | |

| | Distance between Cα atoms (Å) | | | | |
|---|---|---|---|---|---|
| | **R72** | **S193** | **L258** | **T271** | **L273** |
| **R72** | | 15.8 | 10.1 | 5.3 | 6.9 |
| **S193** | | | 20.5 | 17.7 | 13.1 |
| **L258** | | | | 8.6 | 9.7 |
| **T271** | | | | | 5.1 |
| **L273** | | | | | |

**Supplementary Figure 9. Distances between the key functional residues in the crystal structures of a**, AncDHCH1 (pale cyan, PDB entry 6C2C) and **b**, MPH (light orange, PDB entry 1P9E). The panels on the left depict cartoon representations of the crystal structures of the two enzymes, with the five key functional residues highlighted as sticks; because AncDHCH1 lacks a residue at position 193, the nearest residue, *e*192, is used for measurements of inter-residue distances. The panels on the right show the measured distances between the nearest atoms (top) and the Cα (bottom) of the five residues in the crystal structures of the two enzymes. Residues that are close enough for potential physical interactions (distance <4 Å) are bolded.

### Supplementary Figure 10

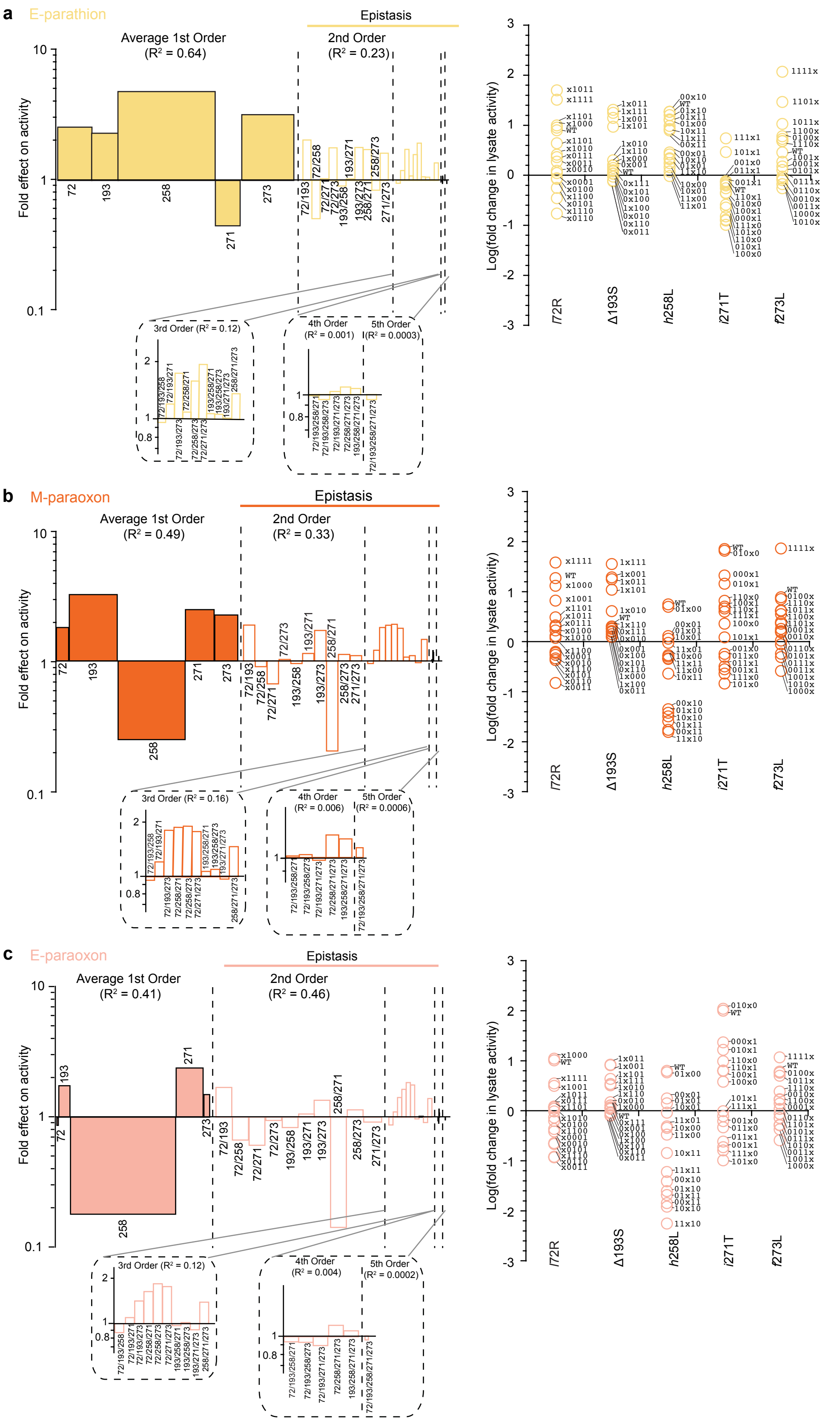


**Supplementary Figure 10. Statistical analyses of singular and epistatic mutational effects for a**, Ethyl-parathion, **b**, Methyl-paraoxon, and **c**, Ethyl-paraoxon. Panels on the left depict the statistical analyses of the mutational effects in the adaptive landscapes. Mutations are indicated by their residue numbers (*i.e.*, 72 denotes the effect of *l*72R; 72/193 denotes the effect of m5+*l*72R/Δ193S). The heights of the bars of the 1st order effects indicate the average effects (fold-change in lysate activity) of the five mutations. Widths of the individual bars in the 1st order effects correspond to the portion of variation in activity ($R^2$ in the linear regression model) attributable to each of the singular effects. The heights of the bars in the 2nd, 3rd, 4th, and 5th order effects correspond to the average epistatic effects of the combined mutations. Widths of the 1st, 2nd, 3rd, 4th, and 5th order effects correspond to the variation attributable to combined mutational effects, calculated as the increase in adjusted $R^2$ of the fit to the data when each term is added to a linear regression model. Panels on the right depict the singular mutational effects of each of the 5 mutations in 16 different genetic backgrounds.

### Supplementary Figure 11

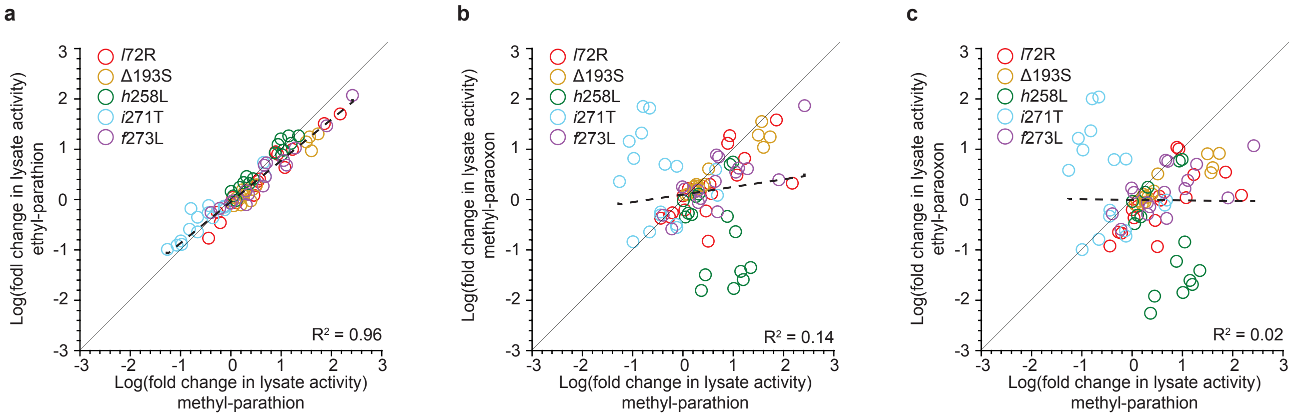


**Supplementary Figure 11. The effects of each of the five key mutations in all 16 possible genetic backgrounds for a**, Ethyl-parathion, **b**, Methyl-paraoxon, and **c**, Ethyl-paraoxon plotted against that for methyl-parathion. The dashed line indicates the linear fit, with the $R^2$ shown in the bottom right corner of each plot. The solid black line running through the graph represent a slope of 1.
